# Joint Speed Discrimination and Augmentation For Prosthesis Feedback

**DOI:** 10.1101/376574

**Authors:** Eric J. Earley, Reva E. Johnson, Levi J. Hargrove, Jon W. Sensinger

**Author notes:** Correspondence should be addressed to E. J. E.

## Abstract

Sensory feedback is critical in fine motor control, learning, and adaptation. However, robotic prosthetic limbs currently lack the feedback segment of the communication loop between user and device. Artificial sensory feedback can close this gap, but sometimes this improvement only persists when users cannot see their prosthesis. suggesting the provided feedback is redundant with vision. Thus, given the choice, users rely on vision over artificial feedback. To effectively augment vision, sensory feedback must provide information that vision cannot provide or provides poorly. Although vision is known to be less precise at estimating speed than position, no work has compared speed precision of biomimetic arm movements. In this study, we investigated the uncertainty of visual speed estimates as defined by different virtual arm movements. We found that uncertainty was greatest for visual estimates of joint speeds, compared to absolute or linear endpoint speeds. Furthermore, this uncertainty increased when the joint reference frame speed varied over time, potentially caused by an overestimation of joint speed. Finally, we demonstrate a joint-based sensory feedback paradigm capable of significantly reducing joint speed uncertainty when paired with vision. Ultimately, this work may lead to improved prosthesis control and capacity for motor learning.

## I. Introduction

When we move our bodies, a complex communication loop is formed between our brains and our extremities. Our brains send efferent commands to our limbs instructing them to move in a specific way. As our limbs carry out these commands, they also send afferent proprioceptive signals back to the brain detailing the positions, speeds, and forces of the limb^1^. From these afferent signals, modifications to the efferent neural drive can correct movement errors and ensure smooth limb control^2^.

Both communication paths are represented by a corresponding internal model. Forward internal models predict future limb movements taking into account the limb’s current configuration and descending signals, while inverse internal models predict the motor command resulting in the limb’s current movement^3^. To develop, adapt, and improve control of the limb over time, these models require knowledge of efferent motor commands (i.e. efference copy) and of the limb’s current configuration and movement (i.e. proprioception). Lack of these proprioceptive signals hampers internal model development and is detrimental to limb control, especially inter-joint coordination^4,5^. Despite its importance in understanding and correcting limb movement, this sense of proprioception is missing for commercially-available robotic prosthetic limbs.

Sensory feedback remains a research priority for prosthesis users^6^. Several feedback methods have been proposed over the past decades, including vibrotactile^7–11^, electrotactile^8^, skin stretch^7^, audio^12–14^ and visual^15^ modalities^16,17^. More complex feedback modalities like peripheral nerve stimulation^18^ and vibration-induced illusory kinesthesia^19^ have also been introduced to great effect.

Sensory feedback typically falls into two categories: tactile feedback of grasping force^9,11,18^, and proprioceptive feedback of limb movement^7,8,10,12–15,19^. Vision is capable of estimating grasping force similarly to tactile feedback^20^, though several grasping force feedback studies have still shown significant benefit to prosthesis control with vision present. However, many proprioceptive feedback studies are conducted with sight of the prosthesis obscured, and the benefit of proprioceptive feedback often diminishes when subjects can see the prosthesis.

During everyday use, prosthesis users visually monitor their device, adopting a distinct gaze pattern. Able-bodied gaze behavior preempts limb movement with eye saccade towards the object of interest^21^, but prosthesis user gaze tends to track the movement of their prosthesis until it reaches the target^22^. This visual monitoring serves to replace the missing proprioception.

When artificial sensory feedback is provided in the presence of vision, the two modalities are integrated according to a weighted sum based on each modality’s uncertainty^23^. Visual estimates of position are highly precise, capable of perceiving changes as small as 1%^24^. In some cases, vision is more precise than even intact proprioception^25^. On the other hand, vision estimates speed with a discrimination threshold of 10%^26^ and a bias towards slower speeds and non-movement (i.e. position)^27^. For either position or speed, if artificial feedback can’t match visual precision, it will be largely ignored in favor of vision. Thus, providing sensory feedback about prosthesis speed should yield a greater benefit than prosthesis position. However, there are several definitions of speed relevant to the movement of a limb.

Limb speed can be defined by the coordinate system (linear speed in Cartesian coordinates, angular speed in polar coordinates) and by the reference frame (absolute speed within a global reference frame, relative speed within a joint-based reference frame). Likewise, feedback provided in joint or global reference frames develop internal models differently, resulting in different generalization to intrinsic or extrinsic error sources^28^. In addition, feedback concerning joint errors is always relevant, but feedback concerning extrinsic errors are only relevant under specific conditions^29^. Despite the importance of joint feedback on tuning the internal models of upper-limb movement, it is not known how precisely vision can perceive joint speed, and thus how effectively artificial proprioceptive feedback can be integrated into such estimates.

The purpose of this study was to investigate visual joint speed perception of biomimetic arm motions, and to determine if these visual joint speed estimates can be augmented with artificial sensory feedback. Subjects observed a virtual two-link arm, analogous to a top-down view of a shoulder, elbow, and hand. Stimuli differed only in the reference frame of interest, and subjects completed two-alternative forced choice tasks to determine just noticeable difference (JND) thresholds. We also tested how joint speed JND varies due to changes in reference frame speed. Finally, we tested a frequency-modulated audio feedback paradigm to evaluate its ability to augment visual speed discrimination.

## II. Methods

### A. Subjects

Experiments were approved by the Northwestern University Institutional Review Board. Methods were carried out in accordance with IRB approval. All subjects provided informed consent before beginning each study. Eight subjects participated in the first and second experiments. Based on a power analysis of simulations using these data, four subjects from the second experiment also participated in the third experiment^30^.

### B. Setup

All protocol and data collection were executed using MATLAB R2017b. Subjects were seated in front of a computer monitor displaying a black two-link system over a uniform white background. The arm had link lengths of 5 cm, widths of 5 points (1.8 mm), and endcap diameters of 6 points (2.1 mm).

Each visual stimulus was presented for 2 seconds, with a 1 second pause between stimuli during which only the white background was shown. Animations were presented at 30 frames per second. Subjects were asked to indicate which stimulus moved faster in the dictated reference frame via a pop-up window prompt. Subjects had unimpaired or corrected vision.

### C. Experiments

Three two-alternative forced choice experiments investigated different aspects of visual speed discrimination. During each experiment, two examples of the two-link arm were displayed to subjects in random order. One stimulus would always move at a nominal speed, whereas the other stimulus would differ from the nominal speed by a magnitude determined by an adaptive staircase. The adaptive staircase was defined as:

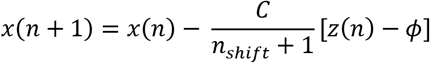

where *x* was the difference in movement speeds between stimuli, *C* was the starting speed difference, *n_shift_* was the number of decision reversals, *ϕ* was the target JND probability (84%), and *z* was a Boolean indicator for the subject’s decision (*z* = 1 when correct and *z* = 0 when incorrect)^31^. Thus, when subjects correctly identified the faster stimulus, the speed difference between stimuli would decrease for the next trial. Likewise, if subjects incorrectly selected the slower stimulus, the speed difference between stimuli would increase for the next trial.

The JND for each condition was calculated as the final stimulus difference *x* tested in the adaptive staircase. The 84% JND has a unique property^32^ in that it is linearly variable with the uncertainty (i.e. standard deviation) of the underlying estimator:

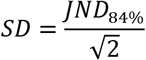

Thus, the 84% JND was converted to uncertainty, normalized, and used as the outcome metric for statistical analyses.

#### 1. Experiment 1: Effect of Speed Type

To determine how discrimination differs between categories of movements, three speed types were tested: *absolute speed, joint speed*, and *linear speed* (Fig. 1). These speed types correspond with different types of proprioceptive feedback that could be provided for prosthetic limbs: speed of a prosthetic joint relative to the torso (*absolute*) or residual limb (*joint*), or speed of the prosthetic end effector (*linear*).

**Figure 1.**
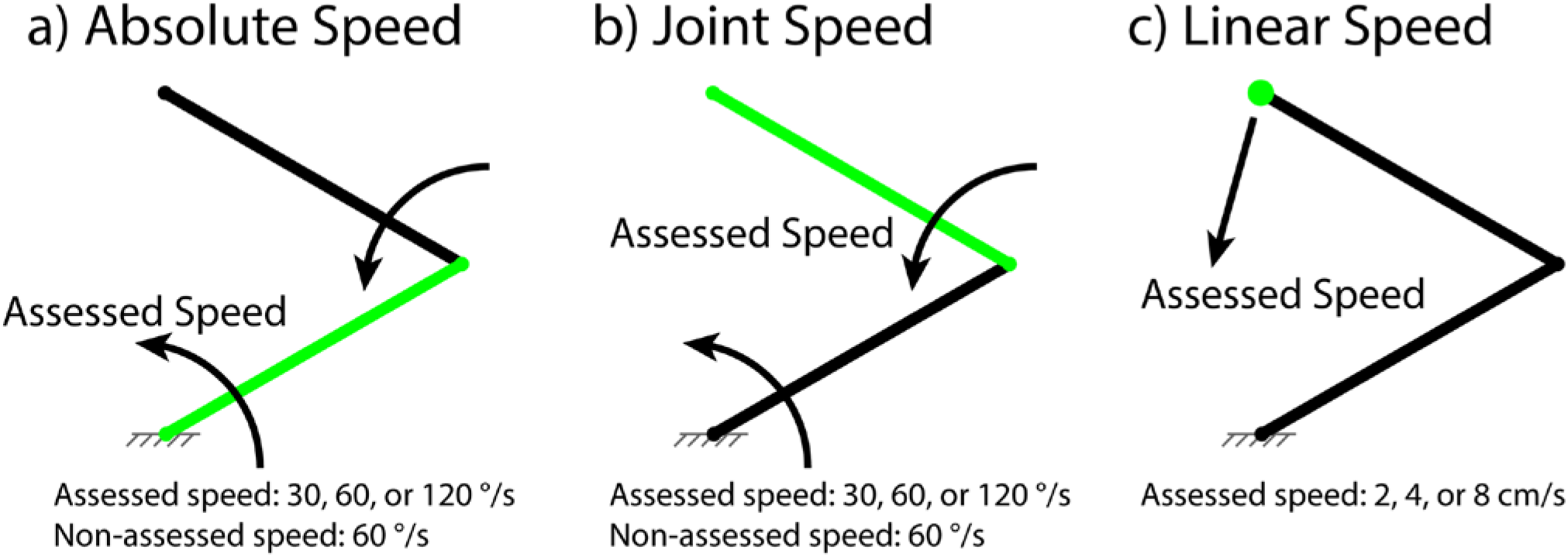
Experiment 1 Setup and Conditions. Assessed component is highlighted green, while all other components are displayed black. Fixture markings in grey are shown here for clarity but were not displayed during experiments. (a) *Absolute speed* condition. Subjects assessed highlighted proximal link speed in three *speed* conditions: 30, 60, or 120 °/s. Distal link rotated at 60 °/s. (b) *Joint speed* condition. Subjects assessed highlighted distal link speed in three *speed* conditions: 30, 60, or 120 °/s. Proximal link rotated at 60 °/s. (c) *Linear speed* condition. Subjects assessed highlighted endpoint speed in three *speed* conditions: 2, 4, or 8 cm/s. Proximal and distal links were driven by endpoint position.

*Absolute speed* refers to rotational movement relative to a global, static reference frame. In this condition, the proximal link moved at a nominal speed of either 30, 60, or 120 °/s counter-clockwise (CCW) for one stimulus, and a speed determined by an adaptive staircase for the other stimulus, starting at *C* = 50%. The distal link moved at a nominal speed of 60 °/s CCW and accelerated and decelerated randomly but equally for both stimuli; thus, the movement profile was not constant, but was identical for both stimuli (Fig. 1a).

*Joint speed* refers to rotational movement relative to a dynamic reference frame, in this case the proximal link. In this condition, the proximal link moved at a nominal speed of 60 °/s CCW and accelerated and decelerated randomly but equally for both stimuli; thus, the movement profile was not constant, but was identical for both stimuli. The distal link moved at a nominal speed of either 30, 60, or 120 °/s CCW for one stimulus, and a speed determined by an adaptive staircase for the other stimulus, starting at *C* = 50% (Fig. 1b).

The random acceleration and deceleration on the proximal link during the *joint speed* condition was implemented to prevent subjects observing absolute speed to estimate joint speed of the distal link by varying the speed of the reference frame. The random acceleration and deceleration on the distal link during the *absolute speed* condition was implemented to match the *joint speed* condition, even though it likely had no effect on estimates.

*Linear speed* refers to movement in a straight line relative to a static Cartesian reference frame. In this condition, the linkage endpoint moved along a straight path at a constant speed of either 2, 4, or 8 cm/s for one stimulus, and a speed determined by an adaptive staircase for the other stimulus, starting at *C* = 50%. The links were driven by inverse kinematics to follow the endpoint (Fig. 1c).

Thus, a total of 9 conditions were tested: 3 speed types, with 3 tested speeds each. Starting positions were randomized for all trials. For *absolute* and *joint speed* trials, the distal link was prevented from crossing the proximal link during movement; invalid starting positions were resampled until conditions were met. Proximal and distal link speeds were bounded between 0 and 180 °/s, preventing clockwise movement and invalid starting positions due to resampling. For *linear speed* trials, the starting position and movement direction were resampled if the endpoint trajectory exceeded the range of the linkage, or if the endpoint didn’t move CCW relative to the origin. The proximal link, distal link, or endpoint were highlighted according to the tested condition.

Statistical analyses performed in RStudio (RStudio, Inc., version 1.1.447) quantified main and interaction effects of the speed type and the observed nominal speed. A Shapiro-Wilk test confirmed normality of the data. A general linear model took the form:

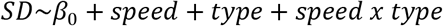

where *speed* was coded as a continuous independent variable in units of octaves (0 at slowest speed, 2 at fastest), and *type* was coded as a categorical independent variable. Because the interaction term was found to be significant, a simple main effects analysis was performed for speed^33^. Corrections for 6 comparisons were made via a Bonferroni correction factor.

#### 2. Experiment 2: Effect of Reference Frame Speed Shift

While the first experiment provided an estimate of *joint speed* perception, it only did so at one reference frame speed. Although results showed a higher uncertainty for *joint speed* observations than for *absolute* or *linear speed* observations, it did not shed any light on possible interaction between changes to the reference frame speed and visual uncertainty. Further, one concern from the first experiment was that during *joint speed* conditions, subjects could conceivably identify the faster joint speed of two stimuli by observing either the joint speed or the absolute rotational speed of the distal link. This ambiguity left open the possibility that that higher uncertainty was due to observing a faster *absolute speed*, rather than due to the *joint speed* nature of the observation itself. We therefore developed a second experiment to determine how joint speed discrimination differs due to changes in reference frame speed. This experiment investigates visual perception of a prosthetic limb while the residual limb is moving non-uniformly. In this experiment, three reference frame conditions were tested. The proximal link rotated at 60 °/s CCW for one stimulus, and a shifted speed of 60, 85, or 120 °/s CCW for the other stimulus; these speeds correspond with an increase of 0, ½, or 1 octave above 60 °/s, respectively. The distal link rotated at 30, 60, or 120 °/s CCW for one stimulus, and a speed determined by an adaptive staircase for the other stimulus, starting at C = 50%. Thus, a total of 9 conditions were tested: 3 reference frame speed shifts, with 3 distal link speeds each (Fig. 1c). Each link was highlighted green at the joint, with a highlight length of 2 cm.

Statistical analyses were performed to quantify how reference frame speed shift magnitude affects uncertainty. A Shapiro-Wilk test confirmed normality of the data. A multiple linear regression model took the form:

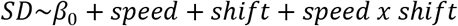

where *speed* and *shift* were coded as continuous independent variables. The interaction term was used to determine if *shift* magnitude impacts uncertainty differently at different *speeds*. The interaction term was not found to be significant (*B* = 0.0002, *t*(68) = 0.304, *p* = 0.762), thus the term was removed and the reduced model was reanalyzed^33^.

After inspecting the data, post-hoc analyses tested the pairs of stimuli subjects choose incorrectly. There were two possible stimulus pairs: one where the speed shift of the reference frame aligns with the faster of the two stimuli, and one where the speed shift occurs with the slower of the two stimuli. The former pair might be considered an easier choice – the correct answer with the faster distal link happens to be the stimulus with the faster proximal link – while the latter pair might be considered a more difficult choice – the correct answer with the faster distal link is the stimulus with the slower proximal link. Therefore, we wanted to determine if *speed* or *shift* impacted the rate of errors due to unaligned stimulus changes (the difficult choice). If there was no impact, subject should make roughly the same number of errors during aligned pairs and unaligned pairs.

Post-hoc statistical analyses were performed using a multiple linear regression model taking the form:

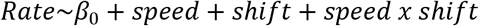

where *speed* and *shift* were coded as continuous independent variables. A Shapiro-Wilk test confirmed normality of the data. The interaction term was not found to be significant (*B* = 0.280, *t*(44) = 1.098, *p* = 0.278), thus the term was removed and the reduced model was reanalyzed^33^.

#### 3. Experiment 3: Effect of Audio Feedback

To determine if joint speed estimates could be improved with supplementary feedback, the *no shift* conditions from the second experiment were repeated. Subjects were provided frequency-modulated audio feedback matching the joint speed of stimuli according to the following equation:

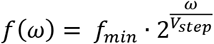

where *f_min_* was the minimum frequency which would be provided when joint speed was zero, and *V_step_* was the speed increase required to increase the audio feedback pitch by one octave. For this study, *f_min_* was set to 220 Hz (A3), and *V_step_* was set to 60 °/s. Audio signals were generated and output with a sampling frequency of 48 kHz. Subjects wore noise-cancelling headphones, and audio was played at a moderate volume. Based on pilot studies, the starting difference *C* between joint speeds was set at 10% to allow the adaptive staircase to converge more smoothly.

Statistical analyses were performed using a general linear model taking the form:

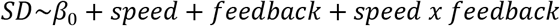

where *speed* was coded as a continuous independent variable and *feedback* was coded as a categorical independent variable. A Shapiro-Wilk test confirmed normality of the data. The purpose of this model was to determine if *vision + audio* improved joint speed discrimination beyond *vision*. The interaction term determined if the benefit of audio feedback was partially dependent on distal link speed, or if benefit was global. A main effects analysis compared *vision* and *vision + audio*. Because the interaction term was significant, and a simple main effects analysis was performed for *speed*^33^. Corrections for 3 comparisons were made via a Bonferroni correction factor.

### D. Data Availability

MATLAB protocol and data analysis code, formatted data files, and R statistical analysis code are freely available for download on the Open Science Framework^34^. Additional data are available upon request.

## III. Results

### A. Experiment 1: Effect of Speed Type

The purpose of experiment 1 was to investigate how visual speed uncertainty differed between *absolute, joint*, and *linear* speed types. In addition, comparing uncertainty across a range of speeds revealed the degree to which uncertainty of each speed type is speed-invariant.

Main effect analysis revealed higher uncertainty for *Joint speed* than either *Absolute* (*t*(36.34) = 4.26, *p* = 0.0008, *d* = 1.23) or *Linear* speeds (*t*(42.79) = 4.24, *p* = 0.0007, *d* = 1.22). *Absolute* and *Linear* speeds were not significantly different (*t*(42.63) = 0.44, *p* > 0.999, *d* = 0.128) (Fig. 3). Thus, our results suggest vision is most uncertain about *joint speed* observations, and thus augmenting joint speed with artificial sensory feedback should yield the greatest improvement in precision.

**Figure 2.**
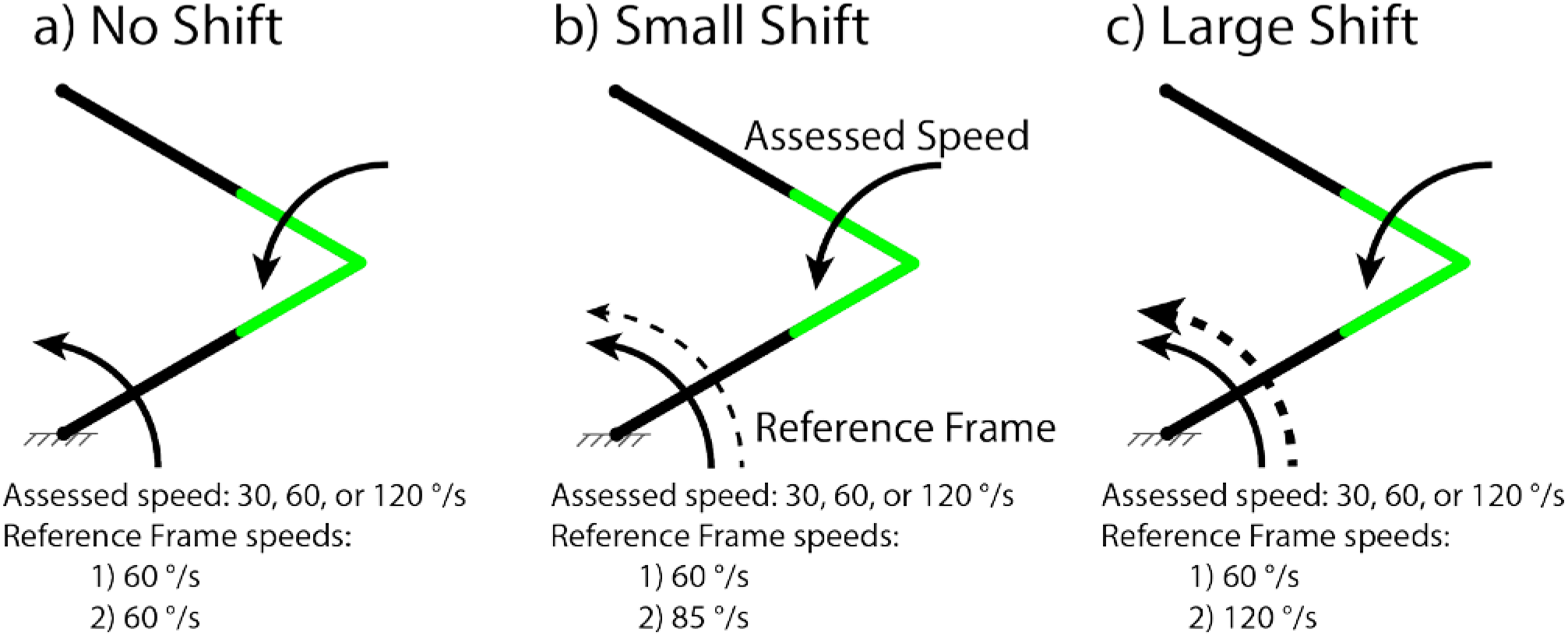
Experiment 2 Setup and Conditions. For each *shift* condition, subjects assessed highlighted joint speed in three *speed* conditions: 30, 60, or 120 °/s. Fixture markings in grey are shown here for clarity but were not displayed during experiments. (a) *No shift* condition. Reference frame rotated at 60 °/s. (b) *Small shift* condition. Reference frame rotated at 60 °/s in one stimulus, and 85 °/s in the other stimulus (c) *Large shift* condition. Reference frame rotated at 60 °/s in one stimulus, and 85 °/s in the other stimulus.

**Figure 3.**
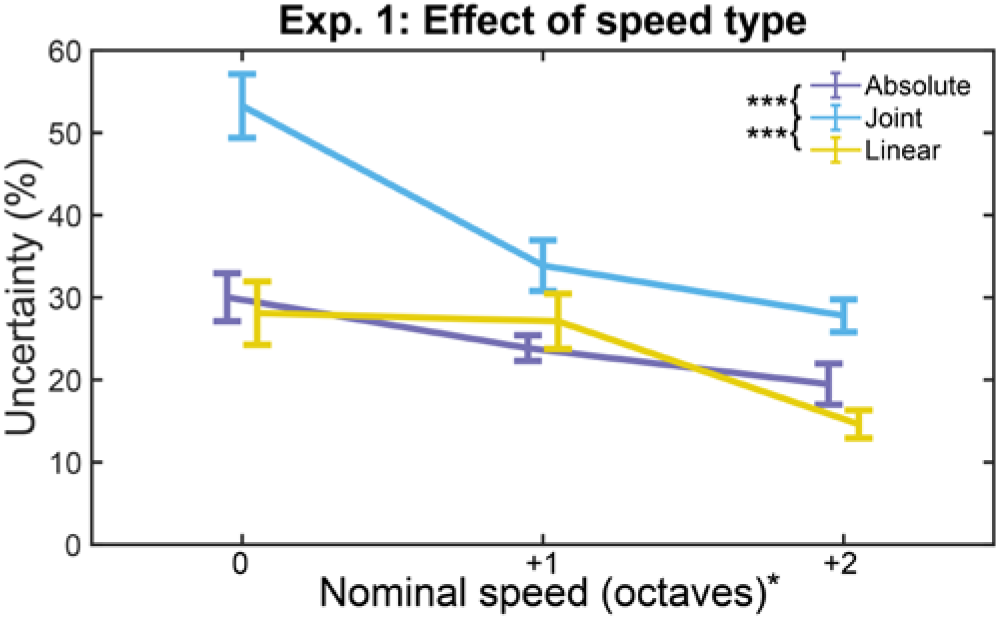
Effect of Speed Type. Uncertainty generally decreases with increasing stimulus speed. Joint speed discrimination is worse than either absolute or linear speed, especially at slow nominal speeds. Error bars show standard error of the mean (SEM). (*)*p* < 0.05, (***)*p* < 0.001

Uncertainty decreased with increasing *speed* for *absolute* (*B* = −0.053, *t*(22) = 3.18, *p* = 0.026), *linear* (*B* = −0.067, *t*(22) = 2.99, *p* = 0.041), and *joint* (*B* = −0.127, *t*(22) = 5.59, *p* < 0.0001) speed types. Significant interaction between *speed* and *type* (*F_4,66_* = 3.60, *p* = 0.033, *η*^2^_*partial*_ = 0.098) suggests that this decrease in uncertainty at higher speeds differs between speed types. This interaction is likely due to the large increase in uncertainty for *joint speed* at low speeds. In particular, in the slowest *joint speed* condition, the assessed joint speed was half as fast as the reference frame speed. Therefore, most of the absolute speed of the distal link was contributed by the proximal link movement, possibly obfuscating the joint speed.

Our results suggest greater uncertainty for visual estimates of joint speed, compared to absolute speed, for biomimetic motions. However, these results alone cannot tell us if this greater uncertainty is due poorer precision of joint speed estimates, or if subjects were estimating the faster absolute speed of the distal link. To remove the confounding factor of being able to estimate joint speed using either method, we followed up with Experiment 2.

### B. Experiment 2: Effect of Reference Frame Speed Shift

Experiment 2 expands upon the *joint speed* results from Experiment 1 by exploring the effect of reference frame speed *shift* on joint *speed* uncertainty. Thus, subjects were unable to make joint speed estimates by observing only the absolute speed of the distal link and were required to consider the speed of the proximal link serving as the moving reference frame.

Main effects analysis showed that both *speed* (*B* = −0.0024, *t*(69) = 9.02, *p* < 0.0001) and *shift* (*B* = 0.0773, *t*(69) = 3.12, *p* = 0.0026) significantly affected uncertainty (Fig 4a). The change in uncertainty associated with changing joint speed confirms the results from Experiment 1 showing similar trends. Additionally, the increase in uncertainty resulting from increased reference frame speed shifts suggests that vision cannot completely filter out reference frame movement during joint speed observations and provides further evidence that joint speed estimates are more uncertain than absolute speed estimates.

**Figure 4.**
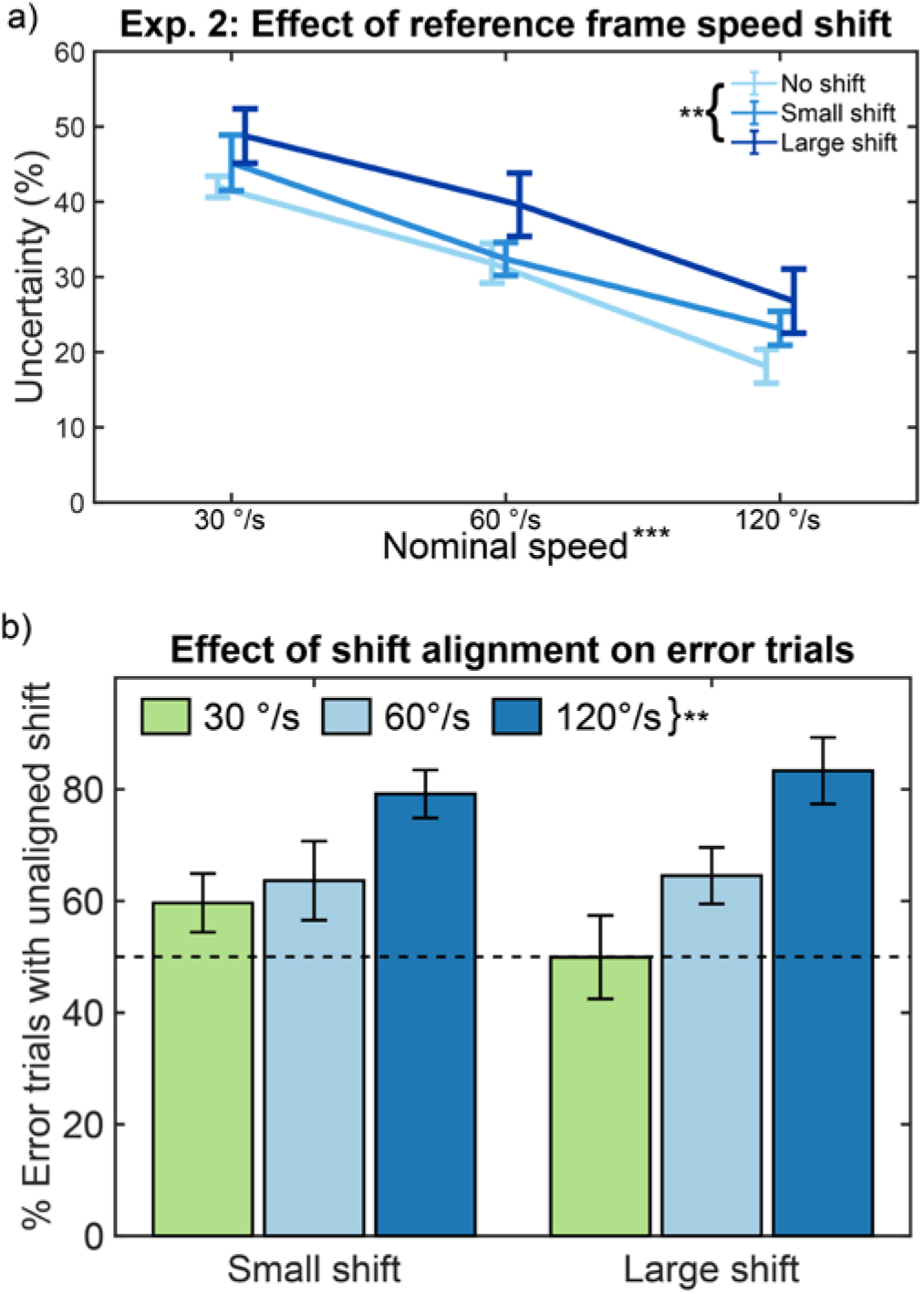
Effect of Reference Frame Speed Shift. (a) Psychophysics results show that increased proximal link speed shift led to increased uncertainty. Error bars show SEM. (b) Shift and increasing joint speed results in an overestimation in joint speed, as shown by an increase in error rate of unaligned trials. The dashed horizontal line indicates the ideal ratio of aligned-to unaligned trials. Error bars show SEM. (**)*p* < 0.01, (***)*p* < 0.001

Post-hoc analyses investigated if either *speed* or *shift* affected the proportion of incorrect stimulus selections where the selected joint speed was slower, but the reference frame moved faster, than the correct stimulus. This rate increased significantly during trials with higher joint *speed* (*B* = 0.293, *t*(45) = 4.59, *p* < 0.0001), but was not affected by *shift* magnitude (*B* = −3.120, *t*(45) = 0.33, *p* = 0.746) (Fig. 4b). This result provides further evidence that vision cannot completely ignore reference frame movement during joint speed observations; instead, reference frame movement may result in an overestimation of, especially, faster joint speeds.

Our results suggest vision cannot completely account for the effect of a moving reference frame when making joint speed estimates, and that a moving reference frame may result in overestimation of the joint speed. Having shown that uncertainty of visual joint speed estimates is greater than absolute speed estimates, we move on to Experiment 3 to determine if vision can be augmented with artificial sensory feedback.

### C. Experiment 3: Effect of Augmentation

Experiment 3 served as a proof-of-concept to show that visual perception of joint speed could be significantly improved with audio feedback. The procedure for Experiment 3 was the same as that for Experiment 2, but subjects wore noise-canceling headphones playing frequency-modulated audio feedback proportional to the speed of the distal joint (i.e. joint speed). Subjects combined visual and auditory cues to arrive at a single joint speed estimate. Main effects analysis revealed significant improvement in uncertainty with audio feedback, over vision alone (*t*(11.06) = 8.14, *p* < 0.0001, *d* = 3.32), providing clear evidence of visual augmentation (Fig. 5).

**Figure 5.**
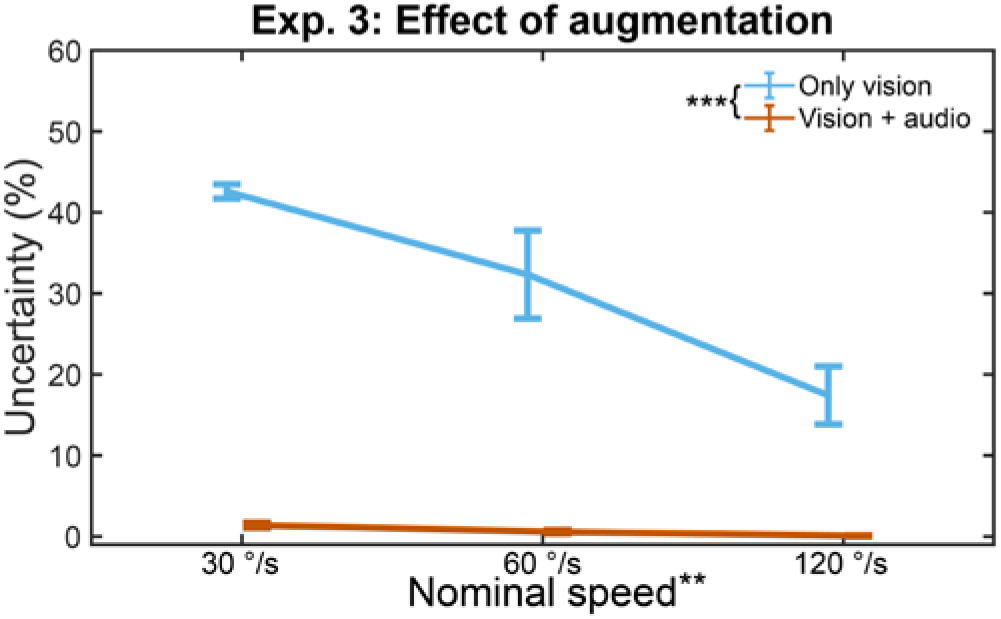
Effect of Augmentation. Visual perception of joint speed is dependent on speed, but audio feedback perception of joint speed is largely speed invariant. Error bars show SEM. (**) *p* < 0.01, (***) *p* < 0.001

Simple main effects analysis revealed speed-varying uncertainty for both *vision* (*B* = −0.0028, *t*(10) = 4.93, *p* = 0.0018) and *vision* + *audio* (*B* = −0.0001, *t*(10) = 3.86, *p* = 0.0094). In addition, interaction between *feedback* and *speed* (*F_1,23_* = 21.85, *p* = 0.0001, *η*^2^_*partial*_ = 0.522) suggests these main effects differ between conditions, particularly that joint speed perception with *vision + audio* is more speed-invariant than joint speed perception with only *vision*. Overall, our results suggest our audio feedback paradigm is sufficient to augment vision when estimating joint speed.

## IV. Discussion

In this study, we investigated visual speed perception of biomimetic arm motions to gain insights for providing sensory feedback for prosthetic limbs. In the first experiment, our results showed that discrimination of linear or absolute speed is between 20% and 30%, whereas discrimination of joint speed is between 30% and 60% (Fig. 3). In the context of providing feedback for prosthetic limbs, our results suggest that providing joint speed feedback will yield the largest improvement to artificial proprioception when users are also able to see the prosthesis.

In the second experiment, our results revealed that variations in the speed of the reference frame reduced discriminatory ability of joint speed observations (Fig. 4a). In post-hoc analyses, we also determined that subjects became more likely to perceive a slower joint speed in a faster reference frame as a faster joint speed, resulting in more incorrect selections (Fig. 4b). This may suggest a multiplicative effect of reference frame speed on joint speed perception. While we found no significant interaction effect between *joint speed* and *shift*, interaction between the two may have plateaued below the magnitudes tested. It is possible that this multiplicative effect arises at smaller reference frame speed shift magnitudes, but further experiments would be required to show this effect. In the context of providing feedback for prosthetic limbs, this second experiment provides evidence that visual joint speed perception is more variable when moving within a time-varying reference frame, such as a prosthetic hand and wrist moving relative to a user’s biological shoulder and elbow. As such, providing joint speed feedback should be most beneficial during tasks requiring coordinated synchronous movement of both robotic and biological joints.

In the third experiment, we showed that visual joint speed estimates can be successfully supplemented with artificial sensory feedback. We provided subjects with frequency-modulated audio cues encoding the speed of the distal link. By playing this joint speed feedback alongside the visual stimuli, joint speed discrimination was reduced below 1%. Additionally, whereas visual discriminatory power varied across nominal speeds, joint speed discrimination was largely invariant with joint speed changes when audio feedback was provided (Fig. 5). This experiment was conducted with no shift to the proximal link speed, the condition with the greatest visual joint speed perception. Because audio feedback is dependent solely on joint speed, proximal link speed shifts which negatively affect visual perception would have no effect on audio perception. Thus, joint speed feedback would provide greater benefits during tasks requiring inter-joint coordination.

Taken together, these results suggest joint speed audio feedback may improve the sense of proprioception for prosthesis users, even when the prosthetic limb is still visible and especially while the residual limb is in motion. This strengthened sensory feedback should, in turn, strengthen internal models associated with reaching tasks, resulting in improved motor learning and control^3^.

Because sensory feedback is merged inversely proportional to each modality’s uncertainty^23^, sensory feedback encoding position will likely not significantly augment proprioception of a prosthetic limb unless it matches or exceeds vision’s 1% uncertainty^24^ or encodes information in a novel way, such as tactile sensation^11,18^ or discrete events in grasping^9^. However, our study suggests that sensory feedback encoding prosthetic joint speed may more significantly augment proprioception of a prosthetic limb due to higher uncertainty in visual estimates of limb joint speed. Additionally, sensory feedback provided for intrinsic joint coordinates should always be relevant to limb control, as opposed to feedback provided in extrinsic coordinates, which may only be conditionally relevant^29^. This persistent relevance would ensure greater generalizability to novel tasks during motor learning with a prosthetic limb^28^.

The major limitation of our work is that all speed estimates were made in a controlled environment: only the two-arm link was shown on screen over a uniform white background, and subjects wore noise-canceling headphones during audio feedback trials. Subjects were exposed to none of the distractions nor divided attention that occur with daily prosthesis use. Additionally, subjects were not asked to control the simulated limb while assessing joint speed, and subjects were able to devote their full attention to visual estimates. Thus, showing that the audio feedback *can* be incorporated into speed estimates does not necessarily mean that the information *will* be incorporated meaningfully during user-in-the-loop control tasks. Prosthesis users typically visually track their prosthesis while in use until they reach an object of interest, at which point visual attention is shared between the object and the prosthesis end effector^22^, but there is no guarantee that prosthesis users with sensory feedback would revert to able-bodied eye gaze behavior^21^. To address this limitation, future real-time experiments will determine the added benefit of joint feedback during reaching tasks.

A limitation of Experiment 2 is that only positive reference frame shifts were tested (Fig. 2). The purpose of this experiment was to remove the possibility that subjects were approximating joint speed by estimating absolute speed of the distal link. Shifts slowing down the reference frame would make it easier for subjects to approximate joint speed with absolute speed estimates, so we opted to only test shifts increasing the reference frame speed. To more rigorously quantify psychophysical measures and the effect of reference frame shifts as a confounding factor, a fully-blocked design with different nominal reference frame speeds, joint speeds, and shift magnitudes and directions would be required.

Audio feedback provides a best-case scenario for augmenting joint speed discrimination. Pitch discrimination within a standard piano range (27.5 Hz – 4.2 kHz) is well below 1%^35^. By contrast, our results suggest visual speed discrimination of 20% or more, though previous research has suggested as low as 10%^26^. Although audio frequency was easily scaled to augment visual joint speed discrimination, other feedback modalities may not have the working range and psychophysical precision to significantly augment visual estimates. Future work will develop a framework to computationally determine the minimum feedback range required for artificial sensory feedback to improve biological observations.

## V. Conclusions

Lack of sensory feedback is a major limitation for modern prosthetic limbs. It is important to not only develop artificial sensory feedback for these limbs, but also to strive for feedback that is more than situationally beneficial. To this end, we investigated human visual perception of arm motion to determine its strengths and, particularly, weaknesses. Our work suggests that vision is most uncertain about joint speed observations, and that it is possible to improve these estimates with artificial sensory feedback. Because this feedback improves joint speed perception even in the presence of vision, we anticipate our proposed feedback system improving myoelectric prosthesis control in a variety of daily tasks, ultimately leading to an improved sense of independence and quality of life for upper-limb prosthesis users.

## VII. Acknowledgements

The authors thank all subjects who volunteered to participate in this study. Funding for this research was provided by NSF-NRI 1317379 and NIH grant T32 HD07418.

## VIII. Author Contributions

E. E. prepared and conducted the experiments, performed statistical analysis, and handled data availability. All authors planned the experiments, prepared and reviewed the manuscript, and approved the submitted version.

## IX. Additional Information

### A. Competing Interests

The authors declare that the research was conducted in the absence of any commercial or financial relationships that could be construed as a potential conflict of interest.

L.H. has ownership interest in Coapt LLC., a start-up company that sells myoelectric pattern recognition control systems. No Coapt products were used as part of this research.

